## Supplementary Material, Tables and Figures for "Aberrant non-canonical NF-κB signalling reprograms the epigenome landscape to drive oncogenic transcriptomes in multiple myeloma"

### Supplementary Tables

***Supplementary Table 1. Primers***

| Name | Sequence |
|---|---|
| Cloning | |
| NFKB2Ex6 F | CACCTCCTAGATCTGTAACTACGA |
| NFKB2Ex6 R | AAATCGTAGTTACAGATCTAGGAA |
| RGS1 SE g1F | CACCGAGTTCAAAGGGGATGTCCAG |
| RGS1 SE g1R | AAACCTGGACATCCCCTTTGAACTC |
| RGS1 SE g4F | CACCGCTCACTTTTGAAGTAGATGC |
| RGS1 SE g4R | AAACGCATCTACTTCAAAAGTGAGC |
| BCL2 SE g7F | CACCGGCATATGGCGTATAAACAC |
| BCL2 SE g7R | AAACGTGTTTATACGCCATATGCC |
| BCL2 SE g8F | CACCGAAACCGGACAGGTGCTGAG |
| BCL2 SE g8R | AAACCTCAGCACCTGTCCGGTTTC |
| BCL2 SE g9F | CACCGAAAGATTTCCCCGCACAGTG |
| BCL2 SE g9R | AAACCACTGTGCGGGGAAATCTTTC |
| BCL2 SE g10F | CACCGGACACTGGAGTCTGACTAG |
| BCL2 SE g10R | AAACCTAGTCAGACTCCAGTGTCC |
| BCL2 SE g11F | CACCGAAGGGAAATCAACAGCACGT |
| BCL2 SE g11R | AAACACGTGCTGTTGATTTCCCTTC |
| BCL2 SE g12F | CACCGTTTTCCAAAATGGTACCCTG |
| BCL2 SE g12R | AAACCAGGGTACCATTTTGGAAAAC |
| RGS1 OE - F | CTAGCTAGCATGCGCGCAGCAGCCATCTCCA |
| RGS1 OE - R | CGACCGGTTCACTTTAGGCTATTAGCCTGCA |
| shRGS1#2 F | CCGG ATTGAAAGGAACCACTCATTCCTGCAG GAATGAGTGGTTCCTTTCAATTTTTG |
| shRGS1#2 R | AATTCAAAAA ATTGAAAGGAACCACTCATTCCTGCAG GAATGAGTGGTTCCTTTCAAT |
| shRGS1#4 F | CCGG GCATTCAGATGCTGCTAAACA CTGCAG TGTTTAGCAGCATCTGAATGC TTTTTG |

| | |
|---|---|
| shRGS1#4 R | AATTCAAAAA GCATTCAGATGCTGCTAAACA CTGCAG TGTTTAGCAGCATCTGAATGC |
| GRAP2 Sh g1-F | CCGGGGAGGCAGCCTTGACATAAATCTGCAGATTTATGTCAAGGCTGCCT CCTTTTTG |
| GRAP2 Sh g1-R | AATTCAAAAAGGAGGCAGCCTTGACATAAATCTGCAGATTTATGTCAAGGC TGCCTCC |
| GRAP2 Sh g2 - F | CCGGGCGAGACAACAAGGGTAATTACTGCAGTAATTACCCTTGTTGTCTC GCTTTTTG |
| GRAP2 Sh g2 - R | AATTCAAAAAGCGAGACAACAAGGGTAATTACTGCAGTAATTACCCTTGTT GTCTCGC |
| Genotyping | |
| RGS1SE F | TTTGCCAAACATGCAGAGTC |
| RGS1SE R | TTTGGCAACAAAACCCTTTC |
| BCL2SE1 F | TTTCTGTACCCCAGGAGGTG |
| BCL2SE4 F | CTCTTGGGCTGTTTTTCCAA |
| BCL2SE2 F | GGAAGACCTGCCAGAGTGAG |
| BCL2SE3 R | CGGCCACCAGGTAAAAAGTA |
| BCL2SE4 R | GAAGAGGGGACTCTGCACTG |
| BCL2SE2 R | CCCTGTGTAGCAAAGGGAAA |
| BCL2SE3 R | CGGCCACCAGGTAAAAAGTA |

***Supplementary Table 2. Summary of features across concordantly regulated EPI pairs***

| Features | Enhancers (Super-Enhancers) | Interactions | Genes | EPI Pairs |
|---|---|---|---|---|
| Upregulated | 75 (19) | 90 (24) | 77 (20) | 99 (26) |
| Downregulated | 357 (127) | 543 (217) | 351 (116) | 695 (283) |

***Supplementary Table 3. Genes linked to Proximal or Distal SEs***

| Group | Gene | log2(FC) | FDR |
|---|---|---|---|
| Proximal | AL358473.1 | -2.09 | 1.41E-02 |
| Proximal | KIF21B | -1.61 | 6.03E-03 |
| Distal | AC126696.3 | -1.55 | 7.00E-02 |
| Proximal | LAIR1 | -1.28 | 4.90E-06 |
| Distal | STOM | -1.08 | 6.58E-13 |
| Proximal | LINC01686 | -1.02 | 1.01E-02 |
| Distal+Proximal | WIPI1 | -0.97 | 1.78E-02 |
| Distal | LINC02362 | -0.86 | 5.53E-07 |
| Proximal | ERN1 | -0.85 | 5.18E-06 |
| Proximal | ANKRD36BP2 | -0.80 | 1.80E-08 |
| Distal | RHOD | -0.79 | 2.53E-04 |
| Distal+Proximal | TMSB4X | -0.78 | 9.41E-12 |
| Distal+Proximal | RGS16 | -0.77 | 2.58E-07 |
| Distal | IRF2BP2 | -0.67 | 2.74E-07 |
| Distal+Proximal | ADTRP | -0.67 | 2.42E-04 |
| Distal+Proximal | AL022724.3 | -0.66 | 1.90E-03 |
| Proximal | AL360182.2 | -0.65 | 7.58E-02 |
| Distal | AL160408.2 | -0.64 | 3.53E-05 |
| Proximal | UBC | -0.63 | 4.46E-05 |
| Distal | AL365272.1 | -0.57 | 3.54E-04 |
| Proximal | CD48 | -0.51 | 6.00E-05 |
| Distal+Proximal | CREG1 | -0.51 | 2.57E-05 |
| Proximal | MXI1 | -0.46 | 1.53E-03 |
| Distal+Proximal | WWC3 | -0.45 | 1.68E-02 |
| Distal | UBALD2 | -0.42 | 1.52E-02 |
| Distal+Proximal | DUSP22 | -0.38 | 1.50E-02 |
| Distal+Proximal | NDUFAF6 | -0.38 | 3.94E-02 |
| Distal+Proximal | QPCT | -0.38 | 5.38E-03 |
| Proximal | SYNGR2 | -0.35 | 4.26E-02 |
| Proximal | NFIL3 | -0.34 | 8.96E-02 |
| Proximal | SUB1 | -0.34 | 2.94E-02 |
| Distal+Proximal | CYTIP | -0.32 | 3.21E-02 |
| Proximal | GALM | -0.31 | 5.26E-02 |
| Distal | PHF19 | -0.30 | 7.20E-02 |
| Distal | IL6ST | -0.28 | 4.97E-02 |
| Distal+Proximal | SEPTIN6 | -0.28 | 7.16E-02 |
| Distal | FHL1 | -0.26 | 8.89E-02 |

### Supplementary Figures

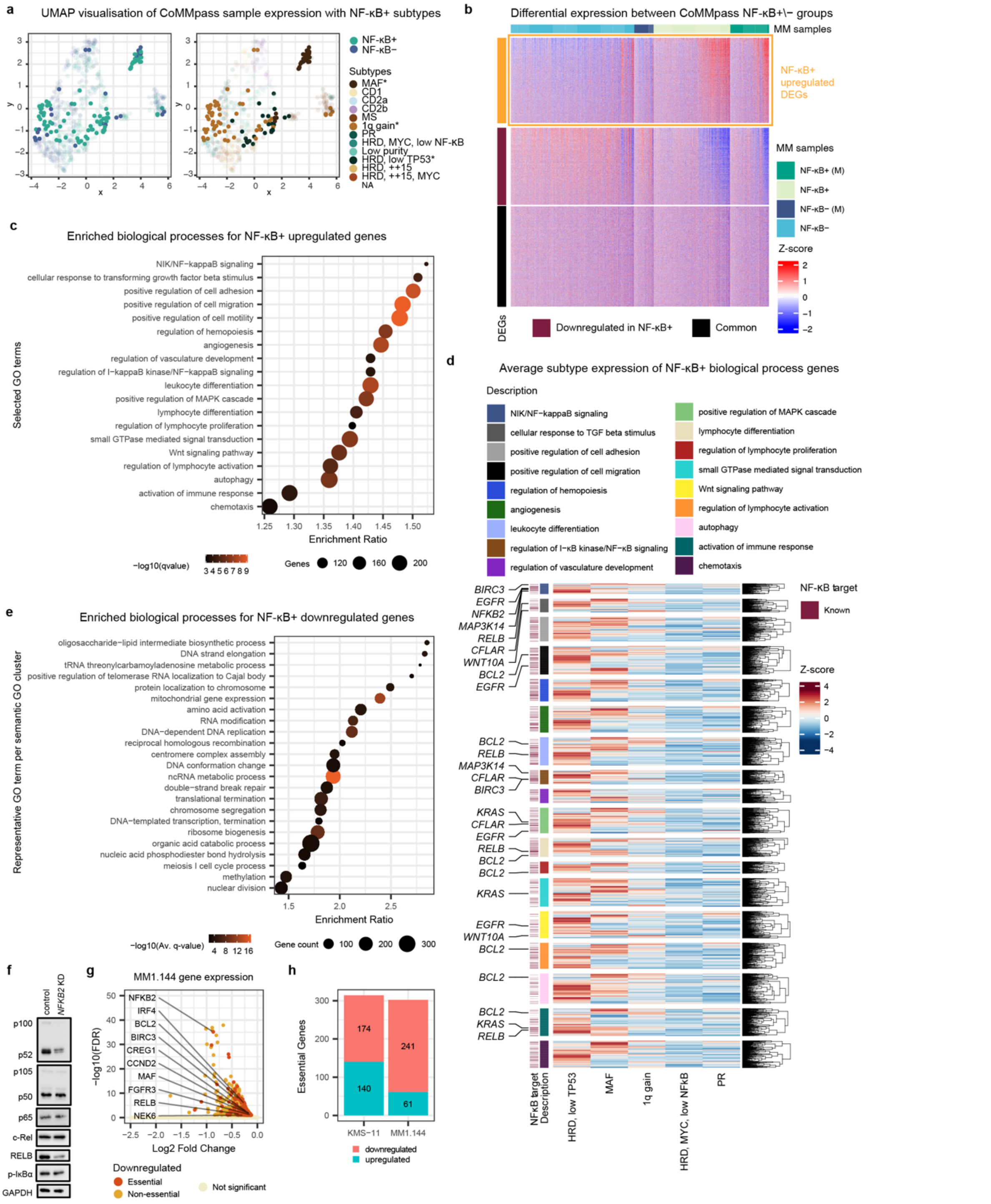


***Supplementary Figure 1. NF-κB+ tumors display gene signatures of 1q gain, HRD low TP53 and MAF subtypes that regulate critical processes in multiple myeloma***
(**a**) UMAP visualisation of CoMMpass sample similarity based on gene expression counts after normalisation and variance stabilising transformation. (**b**) Differential gene expression profiles between NF-κB+ and NF-κB- groups with or without mutations (M), are shown as Z-score of variance stabilised counts for genes with a base mean > 10. (**c**) Gene ontology enrichment analysis (hypergeometric; q-value <= 0.01) showing NF-κB+ samples overexpress genes associated with biological processes often involved with cancer progression. (**d**) Average subtype expression for each gene mapped to selected GO terms identified as enriched in NF-κB+ samples. Known NF-κB targets are indicated. (**e**) Biological

processes significantly enriched (hypergeometric test; Q-value <= 0.01) in NF-κB+ downregulated genes. GO terms were hierarchically clustered based on their semantic similarity with a single representative term chosen for each cluster [1]. Gene count, average enrichment ratio and Q-value for each term/cluster are plotted. (**f**) Western blot showing changes to factors involved in the canonical (NFKB1: p105/p50, p65, c-Rel, p-IκBα) and non-canonical NF-κB (NFKB2: p100/p52, RelB) pathways upon CRISPR-Cas9 knockdown of *NFKB2* in MM1.144. (**g**) Downregulated genes identified following *NFKB2* knockdown in MM1.144, with essential genes highlighted and named. (**h**) Counts of genes deemed essential for multiple myeloma identified as differentially regulated during p52 knock down in KMS-11 and MM1.144.

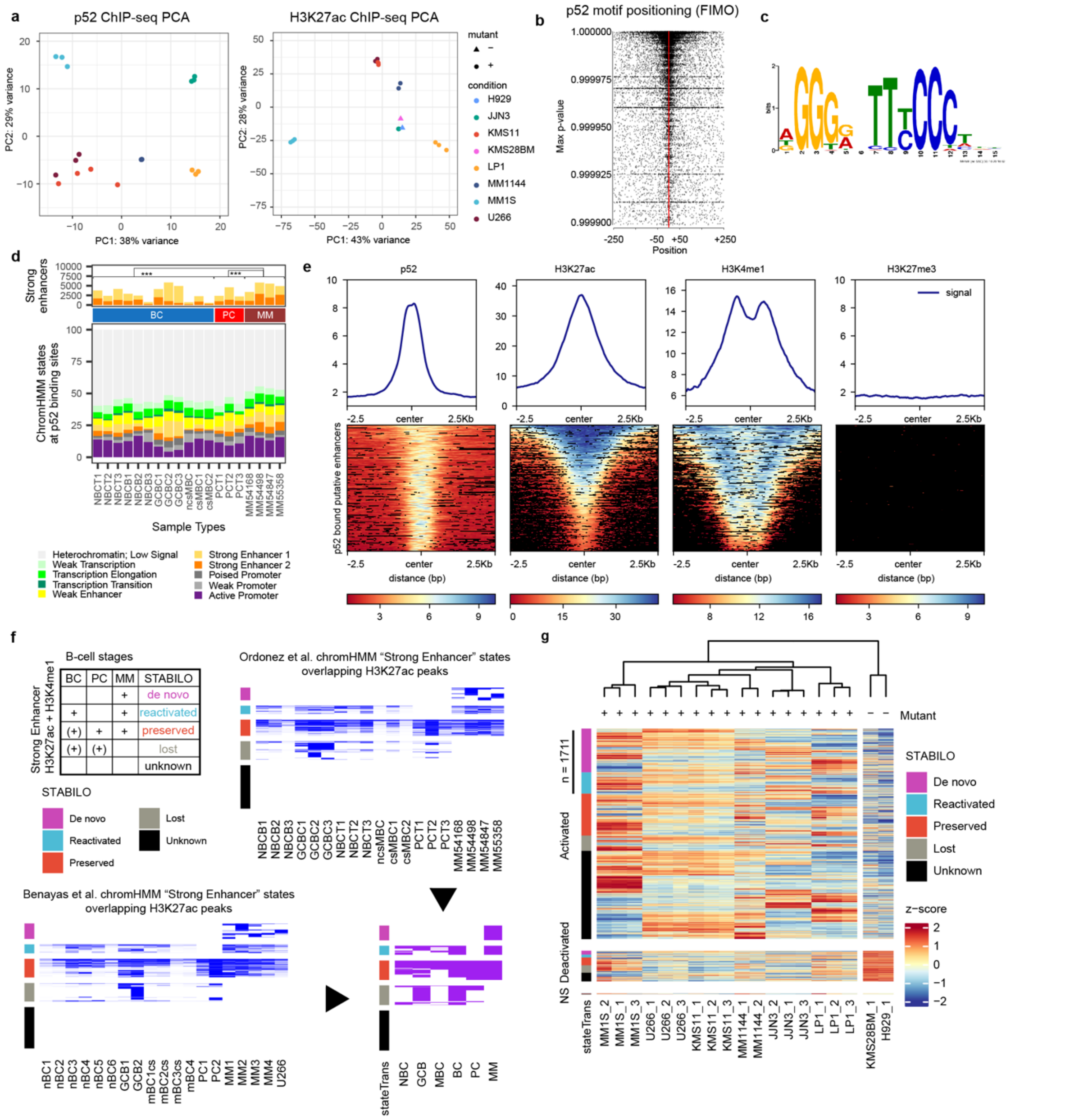


### *Supplementary Figure 2. NF-κB/p52, H3K27ac ChIP-seq and STABILO in MMCLs*

(**a**) Principal component analysis of p52 ChIP-seq and H3K27ac replicate counts in consensus peaks across the multiple myeloma cell lines (MMCLs) panel. (**b**) TFBS-landscape plots showing NF-κB2 motif positioning relative to centre of p52 peaks using FIMO. (**c**) NFKB2 motif found across all p52 ChIP-seq peaks using DREME (**d**) Endogenous p52 binding sites mapped to chromHMM states predicted across three tonsil derived Naïve

B-Cell (NBCT), three blood derived Naïve B-Cell (NBCB), three Germinal Center derived B-Cell (GCBC), one non-class switched Memory B-Cells (ncsMBC), two class-switched Memory B-Cells (csMBC), three tonsil-derived Plasma Cell (PCT) and four multiple myeloma (MM) samples [2]. The proportion of p52 binding sites overlapping strong enhancer 1 and 2 states are significantly greater in MM compared to earlier B-cell stages (Z-Test). (**e**) Profile and heatmap visualisation of the signals obtained from p52, H3K27ac, H3K4me1 and H3K27me3 ChIP-seq at p52 bound enhancers putatively defined as p52 and H3K27ac overlapped peaks in intronic or intergenic regions (**f**) Illustration of the logic and data sources behind the STABILO classification. Two separate studies, Ordonez et al. and Benayas et al. implemented a chromHMM model to segment the genome of different types of B-cells as well as MM samples into 12 epigenomic states. States featuring elevated H3K27ac and H3K4me1 signals were considered as markers of strong enhancer activity. We were therefore able to summarise what segments of the genome transition from one of the 12 states to “Strong Enhancer” states across different sample groups: B-Cell (BC), Plasma Cell (PC) and Multiple Myeloma (MM) using 5 major classes “De novo”, “Reactivated”, “Preserved”, “Lost” and “Unknown” (also see Methods) (**g**) H3K27ac signal (Z-score of normalised rLog counts) at putative enhancers identified across mutant MMCLs bound by p52. Each putative enhancer (rows) is annotated with the STABILO classification (see Methods). Dendrogram for samples is generated by hierarchical clustering.

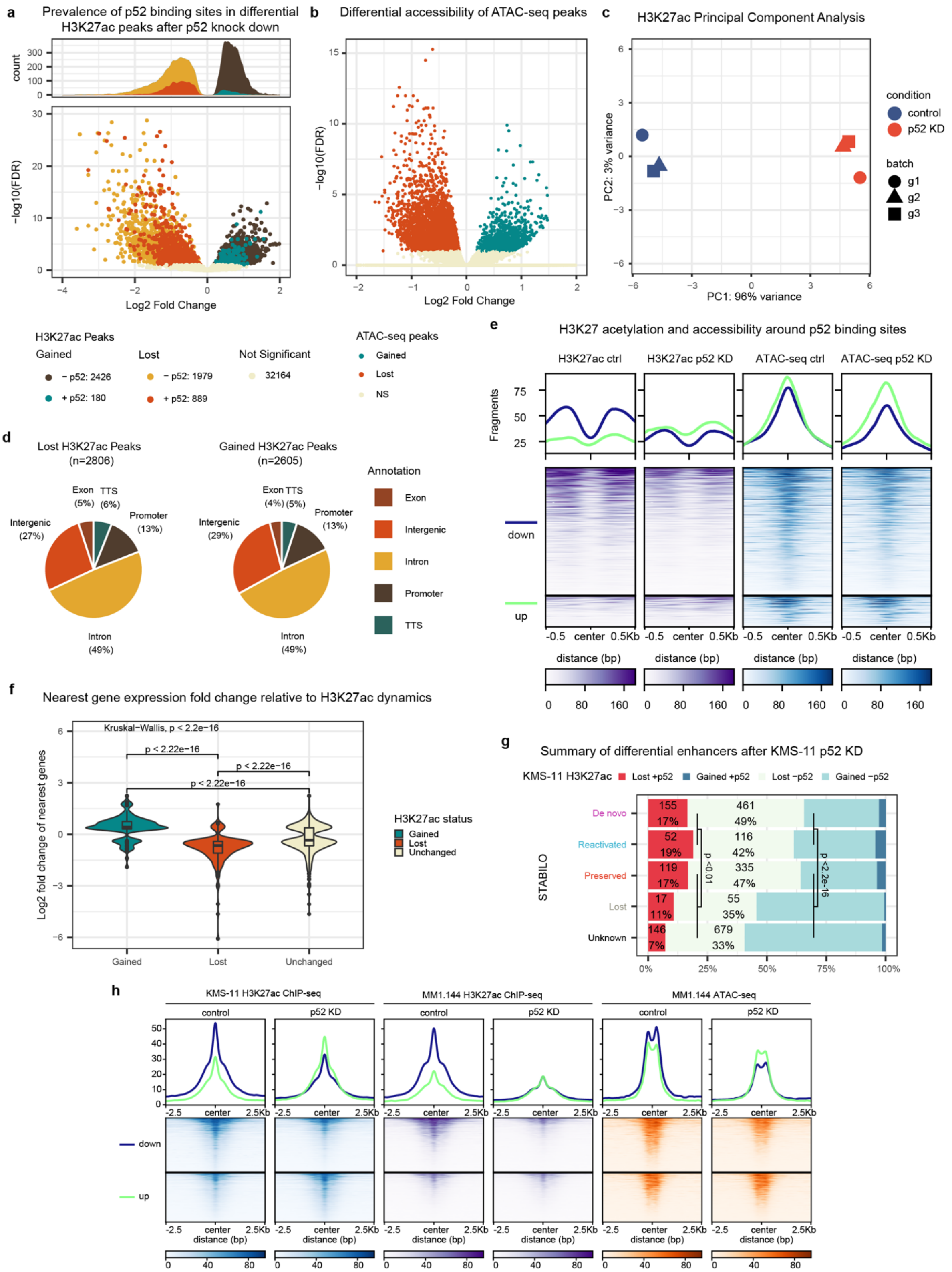


***Supplementary Figure 3. p52 impacts H3K27 acetylation and chromatin accessibility across a diverse collection of enhancers associated with gene expression changes***

(**a**) Loci displaying differential H3K27 acetylation after p52 knockdown (KD) and overlapping p52 binding sites identified in KMS-11 cells. (**b**) Volcano plot of accessibility dynamics at consensus ATAC-seq peaks detected in p52 knockdown (KD) relative to control in KMS-11 cells. (**c**) Principal component analysis of H3K27ac KMS-11 ChIP-seq sample counts in consensus peaks. (**d**) Distribution of differential gained and lost H3K27ac peaks detected in

KMS-11 across genomic features. (**e**) Global H3K27 acetylation signal and accessibility signals plotted as individual profiles and heatmaps centred on p52 peaks. (**f**) Distributions of the significant expression changes for genes nearest to lost, gained or unchanged enhancers identified. Significance between groups was estimated by Kruskal-Wallis and Mann-Whitney U tests. (**g**) Proportions of enhancers bound or unbound by p52 when lost or gained after p52 KD in KMS-11. Dormant (*de novo* + reactivated) enhancers show a significant association with p52-dependent lost enhancers. P-values were calculated using Fisher's Exact Test. (**h**) H3K27 acetylation and accessibility signals obtained in MM1.144 plotted as individual profiles and heatmaps centred on loci exhibiting p52-dependent H3K27 acetylation identified in KMS-11.

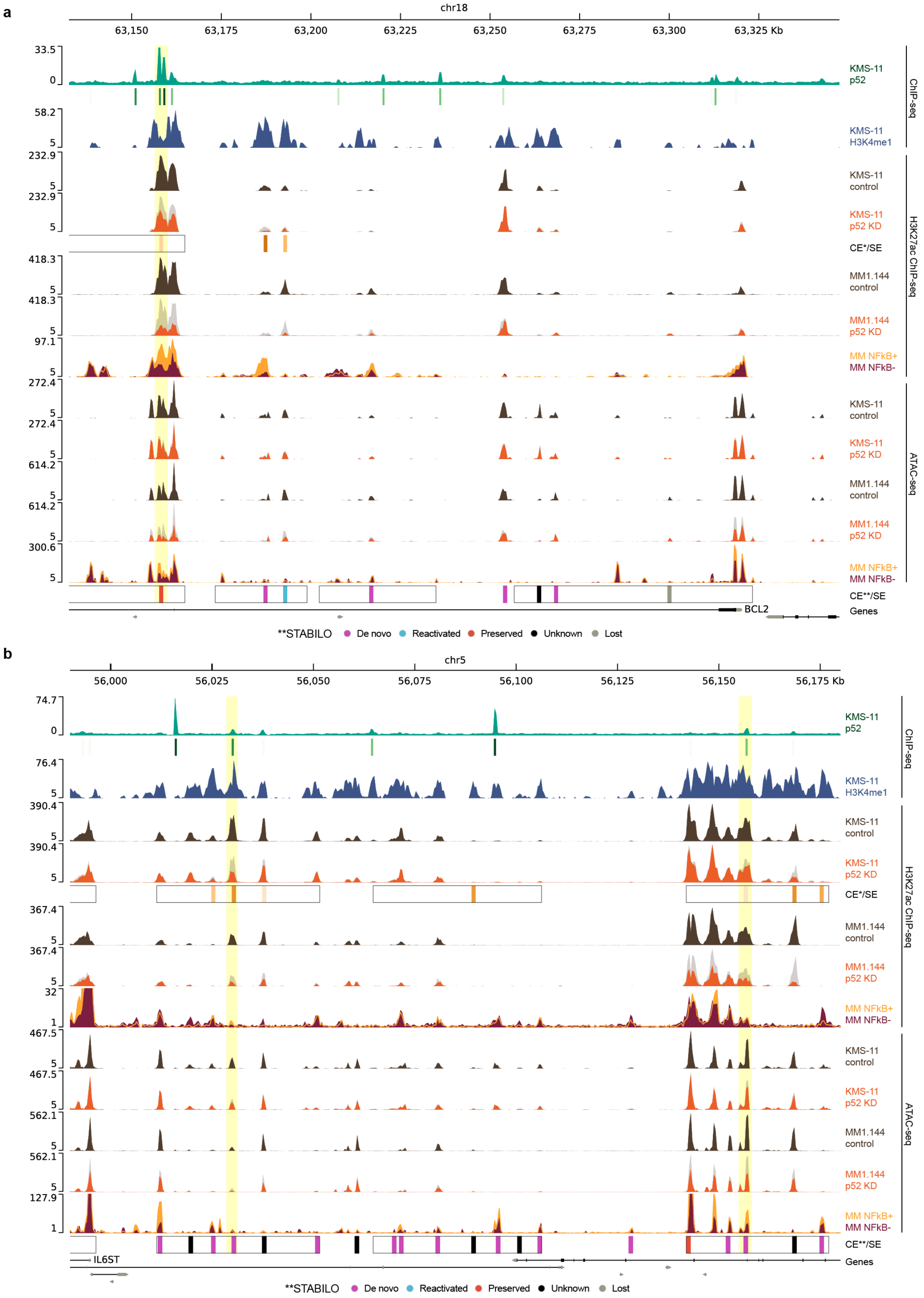


***Supplementary Figure 4. Extended genome browser track visualisations***

Genome browser visualisation for *BCL2* (**a**) and *IL6ST* (**b**) loci encompassing proximal SEs. Tracks display: p52 ChIP-seq signal and peak calls (green) and H3K4me1 ChIP-seq signal obtained in KMS-11; H3K27ac ChIP-seq signal from control and p52 KD experiments in KMS-11 in brown and orange respectively; SEs (black rectangles) and dynamic H3K27ac peaks (orange = lost) called from KMS-11 experiments; H3K27ac ChIP-seq signal from control and p52 KD experiments in MM1.144 (brown/orange); ChIP-seq signal from average of NF-κB+ (yellow; background) or NF-κB- (dark red; forefront) patient samples; ATAC-seq signal from control and p52 KD experiments in KMS-11 (brown/orange); ATAC-seq signal from control and p52 KD experiments in MM1.144 (brown/orange); ATAC-seq signal from average of NF-κB+ (yellow; background) or NF-κB- (dark red; forefront) patient samples; SEs (black rectangles) and constituent H3K27ac peaks classified by STABILO; gene track.

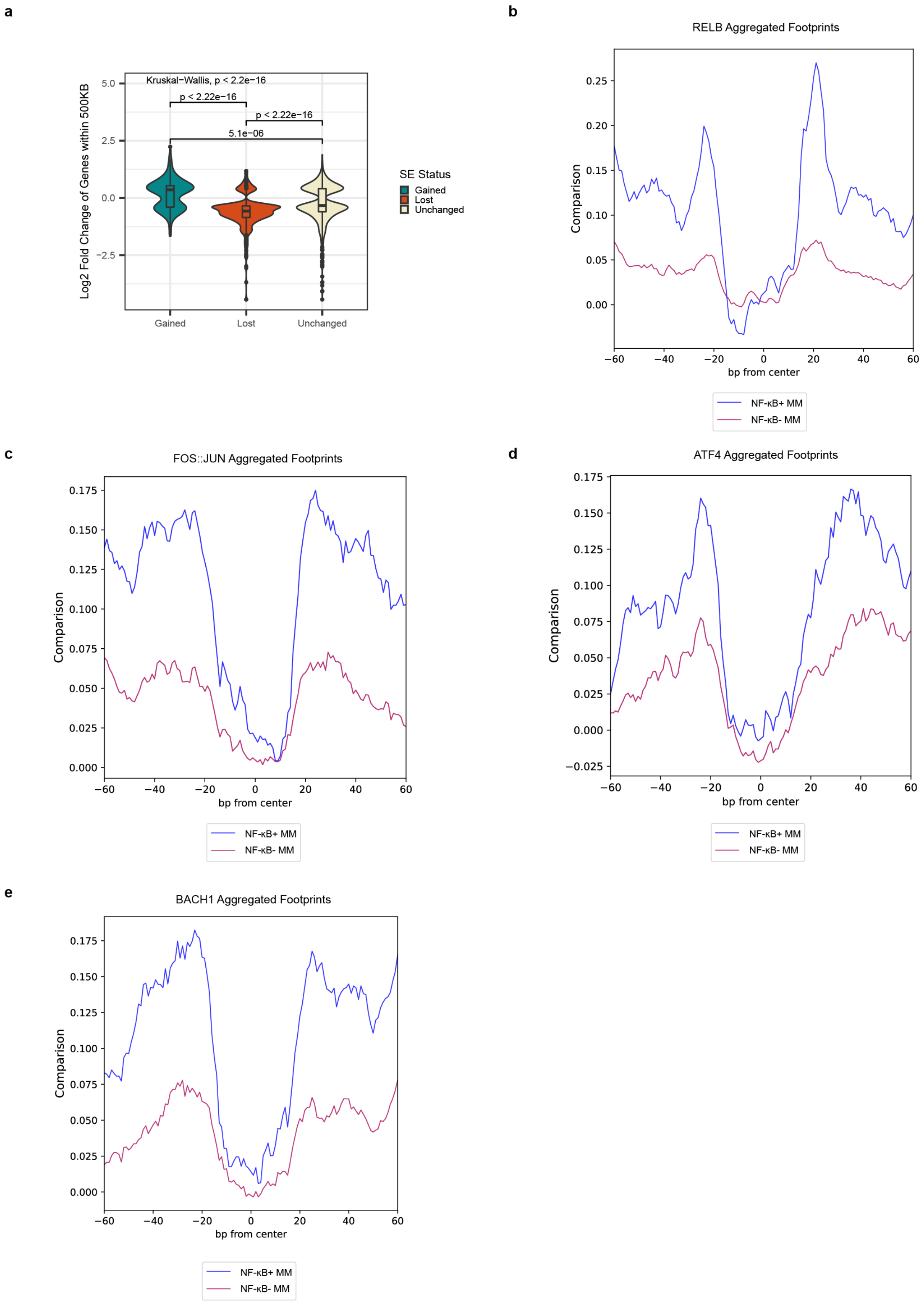


***Supplementary Figure 5. p52 dependent SEs impact proximal gene expression and are bound by several transcription factors enriched in NF-κB+ patients***

(**a**) Significant expression changes for genes found within +/- 500 Kb of lost, gained or unchanged super-enhancers identified following p52 knockdown. Significant differences between groups were tested by Kruskal-Wallis and Mann-Whitney U tests. (**b**-**e**) Aggregated footprints for transcription factors showing differential binding in NF-κB+ MM samples within the SEs identified in KMS-11.

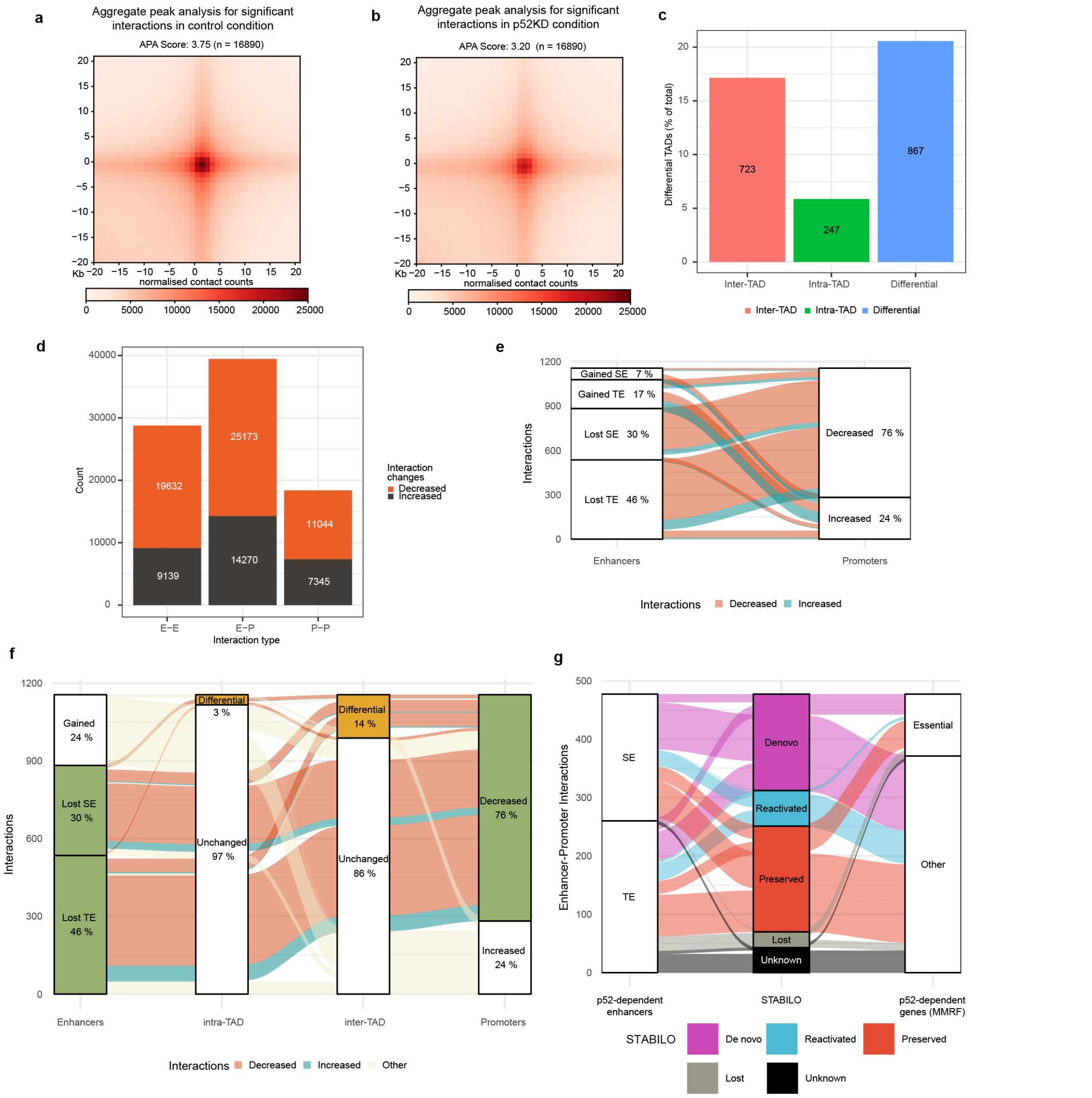


***Supplementary Figure 6. NFKB2 knockdown in KMS-11 cells alters chromatin contact frequency at enhancers***

Aggregate Peak Analysis (APA) of significant loops (**a-b**) called from KMS-11 H3K27ac HiChIP experiments. The p52 KD condition (**b**) showed a reduction in contact frequency at loop peaks compared to control samples (**a**). APA performed on observed/expected transformed 41 x 1 Kb bin matrices with a minimum and maximum range of 50 Kb and 5 Mb respectively. (**c**) Summary of differential TADs detected. 867 differential TADs were detected by hicDifferentialTAD representing 21% of all TADs called. Differential TADs are classified as undergoing inter (723) or intra (247) TAD changes or both (103). (**d**) Numeric breakdown of putative interaction types supported by the loops detected. (**e**) Alluvial plot summarising interaction pairings between differential EP features (Enhancers or Promoters) forming EPI pairs resulting from p52 knockdown in KMS-11 cells. Most downregulated features show downregulation in interactions with their downregulated counterparts. (**f**) Summary of Enhancer to Promoter interactions in the context of TAD changes. Alluvia are coloured to highlight dynamic interactions connecting downregulated enhancer and promoter features (green). As shown by the alluvia, most dynamic EPI pairs occur in unchanged TADs however a minority do occur in TADs undergoing significant intra-TAD or/and inter-TAD changes (yellow). (**g**) Summary of EPIs with concordantly downregulated features (enhancer activity, expression and contacts) upon p52 KD with enriched enhancer or gene activity in NF-κB+ multiple myeloma patients. Enhancer STABILO classification and dependency of target genes are highlighted.

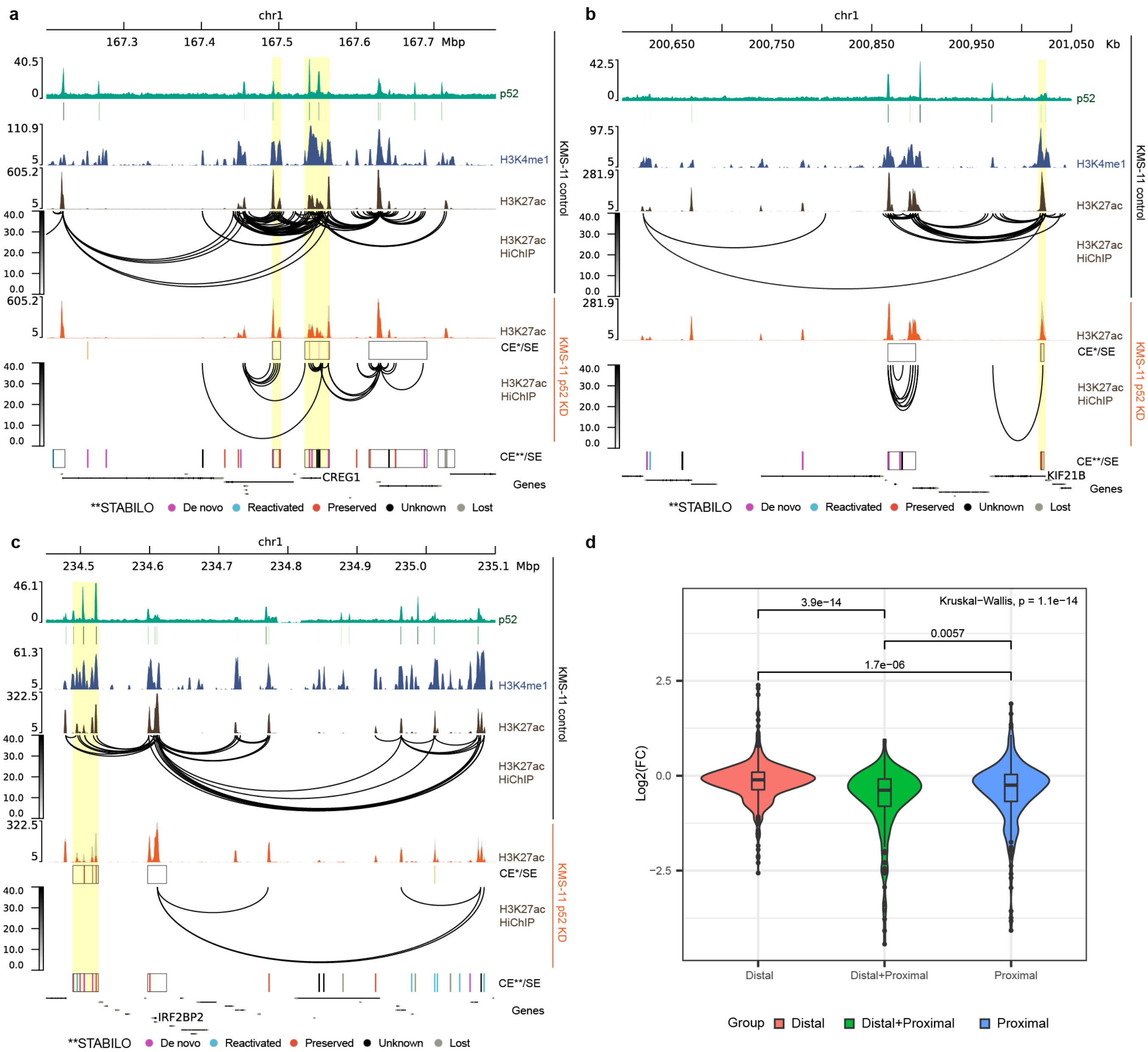


***Supplementary Figure 7. Influence of p52-dependent enhancer distance on gene expression***

Genome browser visualisation for CREG1 (a), KIF21B (b) and IRF2BP2 (c) loci encompassing proximal and distal SEs. Tracks display: p52 ChIP-seq signal and peak calls (green) and H3K4me1 ChIP-seq signal obtained in KMS-11; H3K27ac ChIP-seq signal and HiChIP loops from control (brown) and p52 KD (orange) experiments in KMS-11; SEs (black rectangles) and dynamic H3K27ac peaks (orange = lost) called from KMS-11 experiments; SEs (black rectangles) and H3K27ac peaks classified by STABILO; gene track. All loci feature proximal and distal SEs however distal and proximal enhancers did not show significant changes in H3K27 acetylation upon p52 knock down in KMS-11 in KIF21B and IRF2BP2 respectively. (d) Distributions of the expression changes for genes featuring different combinations of p52-dependent distal and proximal enhancers. Significant differences between groups were tested by Kruskal-Wallis and Mann-Whitney U tests.

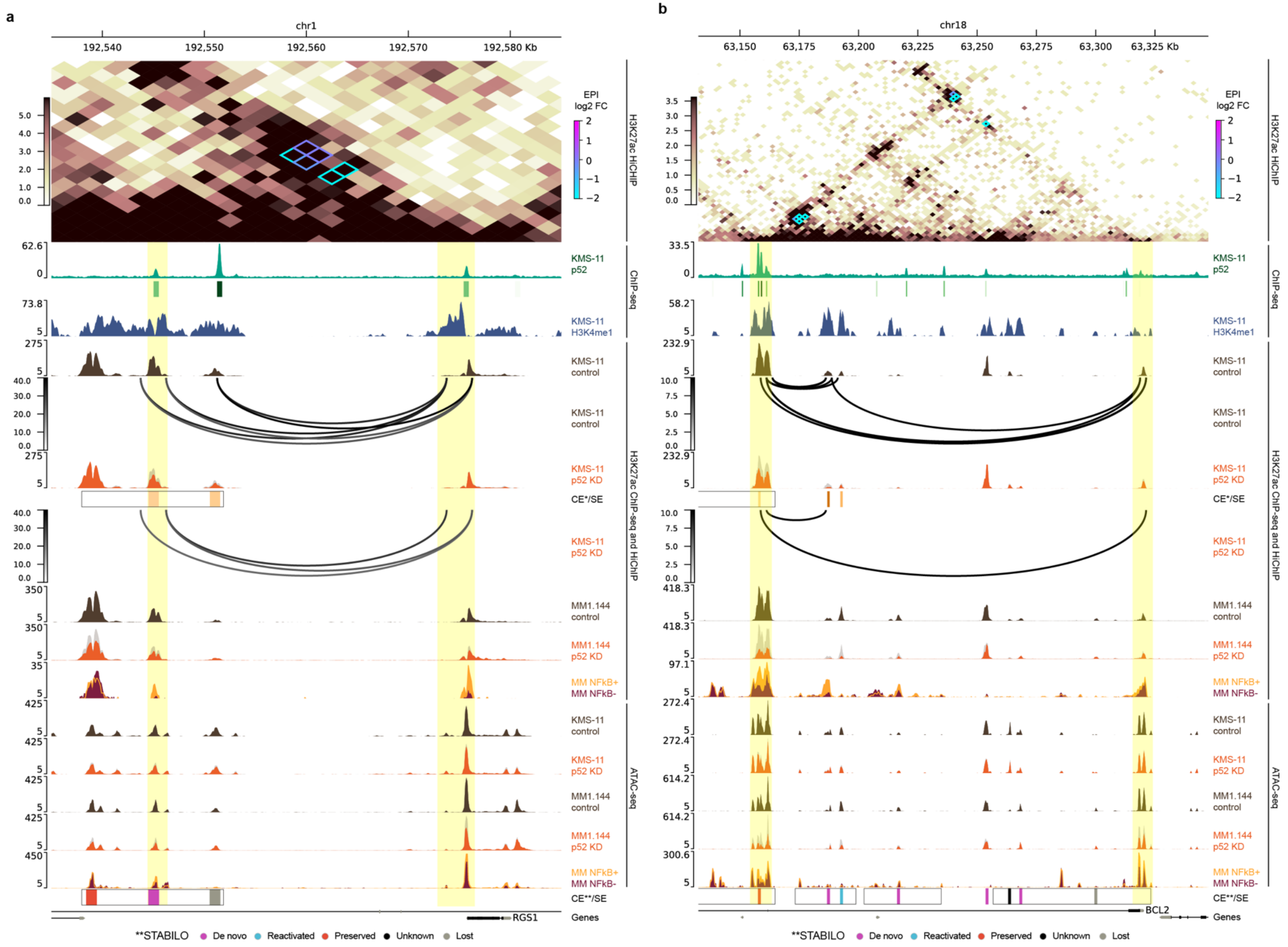


***Supplementary Figure 8. Extended genome browser track visualisations for RGS1 and BCL2***
(**a**) Genome browser visualisation for the *RGS1* and *BCL2* loci encompassing proximal SEs. Tracks display: Contact matrix derived from the H3K27ac HiChIP experiment with differential loops annotated and coloured by log2 fold change; p52 ChIP-seq signal and peak calls (green) and H3K4me1 ChIP-seq signal obtained in KMS-11; H3K27ac ChIP-seq signal and HiChIP loops from control (brown) and p52 KD (orange) experiments in KMS-11; SEs (black rectangles) and dynamic H3K27ac peaks called from KMS-11

experiments; H3K27ac ChIP-seq signal from control and p52 KD experiments in MM1.144 (brown/orange); ChIP-seq signal from average of NF-κB+ (yellow; background) or NF-κB- (dark red; forefront) patient samples; ATAC-seq signal from control and p52 KD experiments in KMS-11 (brown/orange); ATAC-seq signal from control and p52 KD experiments in MM1.144 (brown/orange); ATAC-seq signal from average of NF-κB+ (yellow; background) or NF-κB- (dark red; forefront) patient samples;  SEs (black rectangles) and constituent H3K27ac peaks classified by STABILO; gene track.

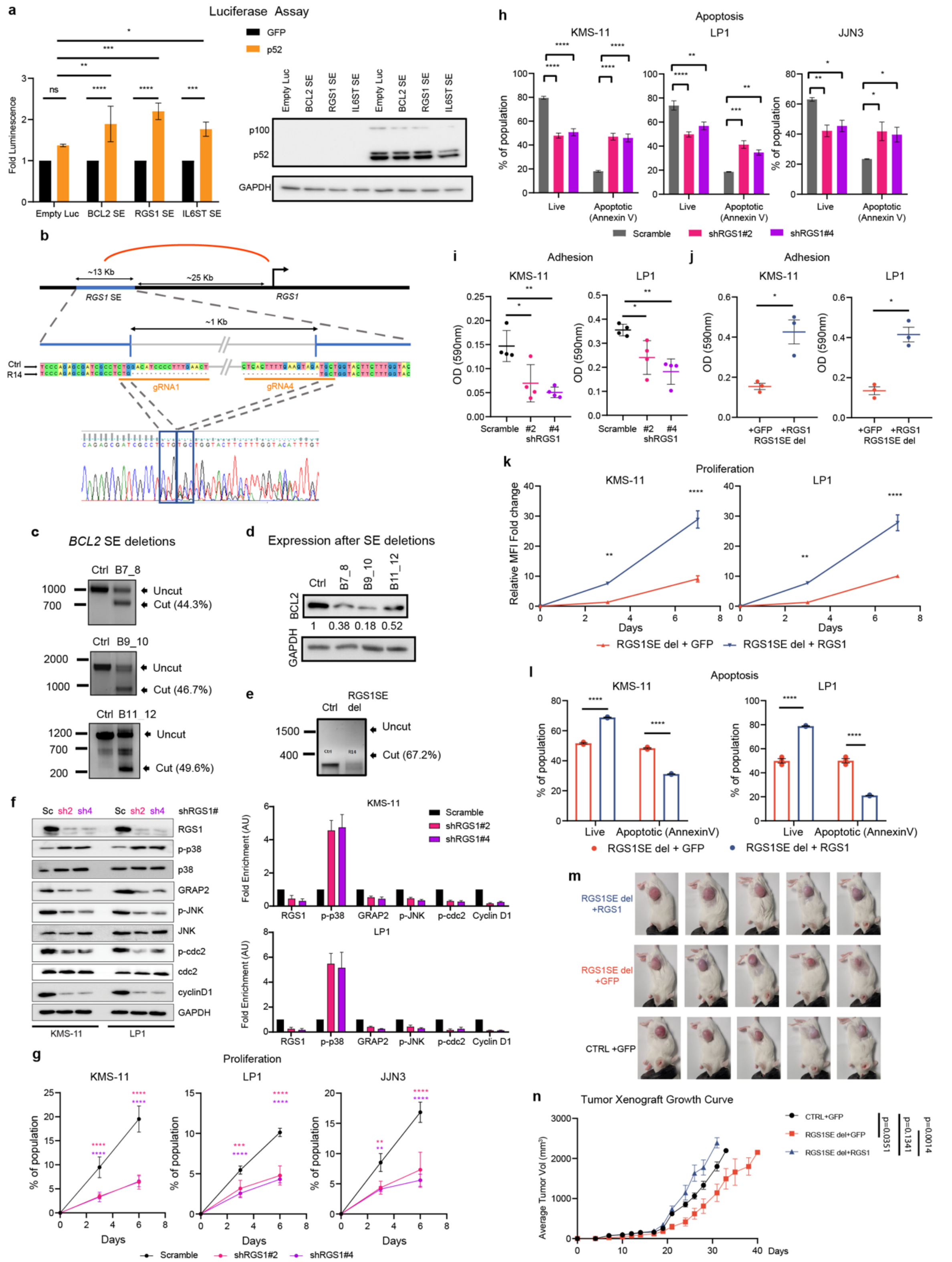


***Supplementary Figure 9. NF-kB/p52 mediated super-enhancer reprogramming impacts the expression and activity of myeloma essential genes***

(**a**) Relative luciferase activity of the p52-bound constituent enhancer of *BCL2*, *RGS1* or *IL6ST* SE regions in 293T cells with or without p52 overexpression. Fold luminescence is calculated by normalising luciferase luminescence reading by renilla luminescence reading. The normalized luciferase activity value in p52 overexpressing cells (fold activation) is then calculated as a fold change to the normalized GFP luciferase activity. Error bars represent mean±SD of three experimental replicates. *P<0.05, **P<0.01, ***P<0.001, ****P<0.0001, ns: not significant. (**b**) CRISPR-Cas9 deletion of a section of the identified super-enhancer region. Illustrated is the SE regulating *RGS1*. gRNA1 and gRNA4 yields a deletion of approximately 1kb within the RGS1 superenhancer. Sanger sequencing of the cut band shows successful deletion of the indicated region. (**c**) Qualitative analysis of *BCL2* enhancer deletion by electrophoresis of genomic DNA PCR products. Wild type genomic DNA (control cells, Ctrl) used as negative controls. B7_8, BCL2SE gRNA7 + gRNA8. B9_10, BCL2SE gRNA9 + gRNA10. B11_12, BCL2SE gRNA11 + gRNA12. Percentage of deletion was calculated from the deletion sample using formula: densitometry (cut band / (cut band + uncut band))%. (**d**) Western blot analysis of *BCL2* protein expression upon SE deletion. Blotting results were evaluated by densitometric analysis, corrected with respect to GAPDH expression and expressed relative to the control (Ctrl). The relative protein amount is reported below the lanes. (**e**) Qualitative analysis of RGS1 enhancer deletion by electrophoresis of genomic DNA PCR products. Wild type genomic DNA (control cells, Ctrl) used as negative control. RGS1SE: RGS1SE gRNA1 + gRNA4. Percentage of deletion was calculated from the deletion sample using formula: densitometry (cut band / (cut band + uncut band))%. (**f**) Effect of *RGS1* knockdown on RGS1 and p38α signalling pathway components in KMS-11 and LP1 cell lines. (**g**) KMS-11, LP1 and JJN3 cells were labelled with CellTrace Blue (#C34574, Invitrogen) and dilution of the dye was tracked via flow cytometry with or without RGS1 shRNA knockdown. MFI (mean fluorescence intensity) fold reduction was calculated relative to day 0. **p<0.01 ***p<0.001, ****p<0.0001; 2-way ANOVA. N=3 (**h**) KMS-11, LP1 and JJN3 cells were transduced with shRNA sequences and after 4 days, they were assessed for apoptosis by FACS. Shown is the percentage of live cells (Annexin V-) and apoptotic cells (Annexin V+). *p<0.05, **p<0.01, ***p<0.001, ****p<0.0001, ns=not significant; 2-way ANOVA. N=3 (**i**) Cell adhesion to fibronectin of KMS-11 and LP1 with RGS1 shRNA knockdown. Points represent the mean OD at 590nm of technical triplicates *p<0.05, **p<0.01 ; 2-way ANOVA. N=4) (**j**) Cell adhesion to fibronectin coated plates. KMS-11 and LP1 RGS1 SE deletion cells overexpressing RGS1. Points represent the mean OD at 590nm of technical triplicates *p < 0.05; t-test. N=3). (**k**) KMS-11 and LP1 with RGS1 SE deletion and RGS1 overexpression were labelled with CellTrace Blue (#C34574, Invitrogen) and dilution of the dye was tracked via flow cytometry. MFI (mean fluorescence intensity) fold reduction was calculated relative to day 0. **p<0.01, ****p<0.0001; 2-way ANOVA. N=3 (**l**) KMS-11 and LP1 with RGS1 SE deletion and RGS1 overexpression assessed for apoptosis by FACS. Shown is the percentage of live cells (Annexin V-) and apoptotic cells (Annexin V+). ****p<0.0001; 2-way ANOVA. N=3. (**m**) Rag-/-, IL2R -/- mice were injected subcutaneously with 5x10^6 KMS-11 cells at the right hind flank. Images taken at D26 post injection. N=5 for each group. (**n**) Average tumor volume of each group measured by electronic caliper over time ± SEM is shown. Mice were sacrificed once tumor volume reached 2000mm$^3$. N =5 for each group.

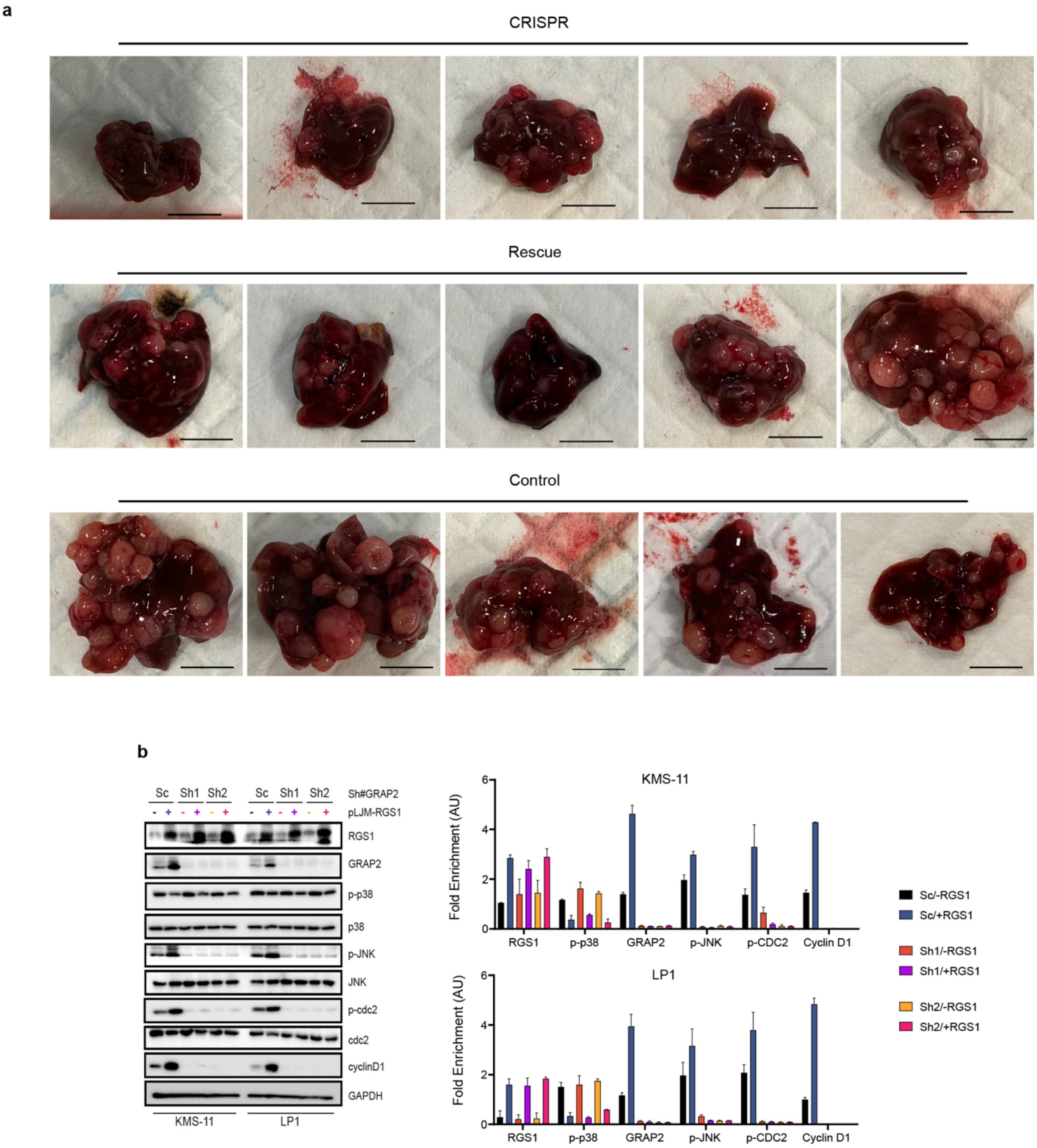


***Supplementary Figure 10. RGS1 SE confers aggressive tumor phenotypes in MM orthotopic models via enhanced expression of RGS1.***
(**a**) Gross anatomy of liver from mice engrafted with *RGS1* SE deleted KMS-11 (CRISPR), *RGS1* SE deleted + RGS1 overexpression KMS-11 (rescue), and KMS-11 (control) cells at the endpoint. Scale bar = 1 cm. (**b**) Representative western blot showing shRNA mediated KD of *GRAP2* attenuates the effect of *RGS1* OE on the activation and expression level of JNK, further downregulating the expression of its target proteins including cdc2 and cyclinD1 in MM cell lines (KMS-11 and LP1). The level of p-p38 remains the same, independent of GRAP2 level, suggesting p38 to be upstream of GRAP2. Bar plots represent the densitometric quantification of the expression levels. Normalization was done taking GAPDH

as loading control and enrichment quantified value (AU) is plotted. N=3 and error bar is SEM.
